# A novel *ex vivo* model of nerve-induced skeletal muscle contraction

**DOI:** 10.1101/2024.10.10.613648

**Authors:** Carole Dabadie, Élise Belaidi, Ioanna Beriatou, Antonio Moretta, Lucas Lemarié, Bruno Allard, Anita Kneppers, Rémi Mounier

## Abstract

Skeletal muscle is a highly organized tissue that possesses the ability to contract and that exerts metabolic, endocrine and paracrine functions. Previous studies explored these functions through *in vitro* models, computational analyses, and endpoint measurements. To include the cellular complexity that modulates muscle endocrine and paracrine functions, and to allow us to study the kinetics of released and/or depleted factors, we developed a novel setup for *ex vivo* contraction of whole skeletal muscle. *Ex vivo* contraction of the *Extensor Digitorum Longus* (EDL) induced by nerve stimulation resulted in a reduction of relative maximal force, an increase in glucose uptake and lactate release, and a depletion of muscle glycogen content, without any overt signs of muscle damage. After cessation of contraction, relative maximal force recovered, and IL6 was secreted. Taken together, we have developed and validated a new model of *ex vivo* contraction that includes the full functional complexity of the muscle and mimics the *in vivo* physiological response to contraction. Importantly, this model supports for the first time to study the kinetics of the endocrine and paracrine response of the muscle to contraction, and to investigate its effects on various cell-types *in vitro*.

## INTRODUCTION

Skeletal muscle is a highly organized tissue, and is predominantly formed by mature contractile cells called myofibers that contain the basic contractile units called sarcomeres. Separated by a dedicated extracellular matrix (ECM) at each structural level, myofibers bundle together to form fascicles that in turn bundle to form the striated skeletal muscle. Finally, fibrous connective tissue or tendons connect muscle to bone to ensure transmission of the contractile forces generated by the muscle to the skeleton. Thus, the architecture of skeletal muscle directly reflects its main function, *i.e.* the generation of force required to maintain posture and drive movement (Frontera & Ochala, 2015; Mukund & Subramaniam, 2020).

While the contractile role of skeletal muscle is well-recognized, skeletal muscle also has critical metabolic functions. For example, skeletal muscle is the major site of glucose uptake in the postprandial state; and as the principal site of insulin-stimulated glucose uptake, it is considered as one of the primary drivers of whole-body insulin resistance (Merz & Thurmond, 2020). These contractile and metabolic roles explain why a loss of skeletal muscle mass and function (called muscle wasting or sarcopenia (Rosenberg, 1997)) is associated with physical frailty and disability (Calvani et al., 2015), increases the risk of metabolic and cardiovascular diseases (Kim & Kim, 2020; Tyrovolas et al., 2020), and even increases morbidity and mortality (de Santana et al., 2021).

Maintenance of skeletal muscle mass and function relies on the intricate relationship between the skeletal muscle contractile and metabolic functions. This relationship is well-exemplified by the release of interleukin 6 (IL-6) from the myofiber upon increased contractile activity and the resulting depletion of substrates such as glycogen (Muñoz-Cánoves et al., 2013). Circulating IL-6 contributes to hepatic glucose production, while IL-6 locally enhances GLUT4 translocation to the plasma membrane to promote glucose uptake into the muscle (Muñoz-Cánoves et al., 2013). Moreover, IL-6 enhances fatty acid oxidation and lipolysis in skeletal muscle *via* the activation of AMP-activated protein kinase (AMPK) (Wolsk et al., 2010), and alters amino acid turnover to support hypertrophic muscle growth (van Hall et al., 2008). Importantly, this demonstrates a coordination of skeletal muscle functions through muscle-derived endocrine, paracrine, and autocrine signaling.

The endocrine role of skeletal muscle and its response to increased contractile activity has been extensively studied (Giudice & Taylor, 2017; Hoffmann & Weigert, 2017). Particularly, the recognition that myofibers produce and release cytokines (called ‘myokines’ (Pedersen et al., 2003)) such as IL-6 into the circulation has sparked studies identifying their role in inter-organ cross-talk (Severinsen & Pedersen, 2020), *e.g.* to redirect substrates to the active myofiber as fuel (Catoire et al., 2014). However, while myofibers are crucial for the key functions of skeletal muscle, they rely on their complex cellular and structural environment for their maintenance and function (Giordani et al., 2019; Rodríguez et al., 2024). Notably, skeletal muscle innervation affects both the quantify and composition of released factors such as myokines (Huang et al., 2024; Rentería et al., 2022). As such, the importance of the skeletal muscle cellular complexity for its endocrine and paracrine role, as well as for its response to increased contractile activity, should be addressed.

Here, we present a novel setup to induce *ex vivo* isometric contraction of whole muscle invoked by electrical pulse stimulation of the nerve. Specifically, we devised a setup that allows the collection of released and extracted factors. This method includes all cell-types and structural components involved in skeletal muscle contraction and mimics the physiological response to acute contraction. Thereby, this setup will help unravel how skeletal muscle mass and function are maintained *via* the complex interplay between cell-types and structural components through paracrine and endocrine signaling upon muscle contraction.

## MATERIALS AND METHODS

### EDL extraction

Mice were housed and maintained in accordance with the French and European legislation in the ‘*Service Commun des Animaleries de Rockefeller*’ (SCAR) animal facility in Lyon (agreement #D693881001). Young adult (8-12 weeks-old) male C57BL/6J mice were anesthetized with isoflurane, and killed by cervical dislocation before muscle extraction. Briefly, the skin of the lower limb was removed, and the lower limb was isolated and placed in Dulbecco’s phosphate-buffered saline (PBS, Gibco #14190169). Then, the *Tibialis Anterior* (TA) was removed and connective tissue overlying the proximal and distal tendon of the *Extensor Digitorum Longus* (EDL) was carefully cut to expose the EDL muscle-tendon complex. Silk sutures (Ethicon #FK889) were tied around the tendons close to the myotendinous junction, and small loops were tied for mounting in the *ex vivo* setup. To extract EDL with the associated nerve, the *Gastrocnemius* muscle was then cut to expose the distal part of the sciatic nerve, and *Peroneus longus* and *Peroneus brevis* muscles were cut to expose the deep peroneal branch of the sciatic nerve innervating the EDL. Connective tissue was removed from the nerve and the nerve was cut 8 to 10mm from the muscle. The EDL, or EDL with the nerve was then extracted by cutting the proximal and distal tendons. Before extraction, *in situ* EDL length was measured between the suture knots (13.95 +/- 0.65mm). Extracted EDL were directly mounted in the *ex vivo* muscle contraction setup.

### Ex vivo nerve stimulation-induced muscle contraction setup

The nerve stimulation (NS) setup is composed of a suction electrode (AMS holder #672440, AMS silver wire #782500, AMS tubing #803400 and Warner capillaries #GC100T-15 coupled with a three-way stopcock and syringe) fixed in a 3D printed micromanipulator (open-source schematic available at https://backyardbrains.com/products/micromanipulator), and a 3D printed customized lid containing two surgical steel pillars that fits a standard 6-well culture plate (**Figure S1A-B**). EDL muscles were mounted at 14mm (*i.e. in vivo* and optimal length), and submerged in pre-warmed (37°C) DMEM high glucose GlutaMAX medium (Gibco #31966021) supplemented with 1% penicillin-streptomycin (Sigma #P4333). The nerve was aspirated into the capillary of the suction electrode, and the mounted EDL with nerve were then acclimatized in a humidified incubator at 37°C and 5% CO_2_ for a 15min period before subsequent force measurements or the electrical pulse stimulation (EPS) protocol.

### Ex vivo muscle force measurement

Muscle force was measured in a separate setup where one surgical steel pillar was replaced by a force-displacement transducer (Grass Instrument Co. #FT03) (**Figure S1C-D**). Maximal isometric force (F_max_) was invoked by 1-2 seconds (sec) stimulation at 100Hz (1 millisecond (ms) pulse width) (**Video S1**). To assess fatigue, EDL muscles were subjected to 3 minutes of repeated 300ms pulse trains at 30Hz (1ms pulse width), with a delay of 700ms between repeats. Force output was recorded in mV using a Powerlab system and Labchart 7.0 software (ADinstruments), and converted to mN or expressed relative to the initial F_max_. All force measurements were conducted at room temperature (20-25°C).

### Induction of ex vivo muscle contraction (EPS protocol)

Repeated muscle contractions to increase the skeletal muscle metabolic demand were invoked by electrical pulse stimulation (EPS) elicited using a constant-current stimulator (Harvard Apparatus 6002 stimulator) controlled by a Powerlab system and Labchart 7.0 software (ADinstruments). EDL muscles were subjected to 4x15min of repeated 360ms pulse trains at 50Hz (resulting in a fused tetanus) (1ms pulse width), with a delay of 29.65sec between repeats (**Figure S1E**). F_max_ was invoked at the start (**Video S1**), and after each 15min series of stimulation (**Figure S1E**). Experiments were conducted in a humidified incubator at 37°C and 5% CO_2_. After completion of the EPS protocol, the muscle and conditioned medium (CM) were snap frozen in liquid nitrogen and stored at -80°C until further analyses. To assess the preservation of force in unstimulated muscles, the non-contracting controls were subjected to F_max_ measurements at the start, and after each 15min interval. For measurements of metabolic responses in muscles and conditioned media (CM) upon nerve-stimulation, non-contracting controls received the EPS protocol stimulation in the absence of a nerve, resulting in no contraction (**Figure S1F**). Moreover, an EPS control CM was generated by subjecting the EPS protocol stimulation to medium in the absence of an EDL (**Figure S1F**).

### Curare treatment

After the first 15min of repeated 360ms pulse trains of the EPS protocol, muscles were treated with 10µM tubocurarine hydrochloride pentahydrate (Curare; Sigma #T2379). After the total loss of force production, curare was washed out by replacing the medium with fresh medium for 3 times.

### Lactate dehydrogenase assay

Lactate dehydrogenase (LDH) activity was determined in CM using a colorimetric enzymatic assay (Theret et al., 2017). Briefly, 8.75μl of supernatant from each sample was transferred to a 96-well plate. 241.25μl of reaction mix (20mM imidazole, 0.05% BSA, 0.28mM NADH) was added to each well, and the absorbance was subsequently measured at 340nm every 15sec during 2min to establish the baseline, and every 15sec during 5min after the addition of 33mM sodium pyruvate. LDH activity was calculated as the consumption of NADH *per* minute *per* μL of sample (mmol/min/μL). Negative activity values were set to 0. Positive control was generated by incubating a mechanically injured EDL in medium for 1 hour (h).

### Creatine kinase assay

Creatine kinase (CK) was measured in CM using a Quantikine mouse CK ELISA kit (R&D Systems #NBP2-75306) according to the manufacturer’s instructions. The concentration of CK was expressed in ng/mL. Negative values were set to 0. Positive control was generated by incubating a mechanically injured EDL in medium for 1h.

### Histological and immunohistochemical analyses

For histological analyses, EDL were embedded in OCR, frozen in liquid nitrogen cooled isopentane, and stored at -80°C until use. Transverse 10-µm-thick cryosections were prepared at -20°C and air-dried at room temperature (RT) for 30min for subsequent hematoxylin-eosin (HE) staining and immunolabeling. For Dystrophin/Laminin immunolabeling, sections were permeabilized for 10min in 0.5% Triton X-100 (Sigma #T8787) and saturated in 2% BSA (Fisher #BP9701) for 1h at RT. Sections were co-labeled overnight at 4°C with primary antibodies directed against Dystrophin (rabbit polyclonal, Abcam #ab 15277, 1:1000) and Laminin [4H8-2] (rat monoclonal, Santa Cruz #SC-59854, 1:1000). Secondary antibodies coupled to Cy5 or FITC (Jackson ImmunoResearch Inc.; 1:200) were incubated for 1h at 37°C. Nuclear counterstain was performed by 10sec incubation with Hoechst 33343 (Sigma #14533; 2µM), and coverslips were mounted with FluoroMount-G mounting medium (Interchim #FP483331). Scanned slides were acquired with a Zeiss Axio Scan.Z1.

### Lactate assay

Lactate abundance was determined in CM using a colorimetric enzymatic assay (Theret et al., 2017). Briefly, 34.5μl from each sample was transferred to a 96-well plate. 34.5μL H_2_0, 8.6μL NAD 60mM and 172.4μL hydrazin buffer (1M glycin, 5mM EDTA, 0.4M hydrazin pH 9) was added to each well. The absorbance was directly measured at 340nm using a microplate reader to establish the baseline, and 30min after the addition of LDH. Lactate concentration was calculated as the production of NADH *per* μL of sample (mmol/μL). Negative values were set to 0.

CM from C2C12 myotubes was recovered as a positive control. To this end, the murine myoblasts C2C12 cell line was cultured in DMEM high glucose GlutaMAX medium (Gibco #31966021) supplemented with 10% fetal bovine serum (FBS, Sigma #F7524) and 1% penicillin-streptomycin (Sigma #P4333). Cells were plated at 30 000 cells *per* cm^2^ onto a Matrigel-coated plate, and differentiated for 9 days in DMEM high glucose GlutaMAX medium (Gibco #31966021) supplemented with 2% horse serum (HS, Sigma #H1270) and 1% penicillin-streptomycin to obtain myotubes. After 9 days, medium was replaced by FBS-free medium, and FBS-free conditioned medium containing lactate was recovered after 24h of incubation.

### Glycogen and glucose assay

Glycogen abundance was measured in EDL muscles using a colorimetric assay. Briefly, muscles were weighed and digested in 30% KOH at 90°C for 20min. Glycogen was precipitated by addition of ethanol (96%) and overnight mixing by inverting the tubes at 4°C. Samples were centrifuged (10min, 8000g, 4°C), and the precipitate was dissolved in 2mL H_2_O. Glycogen was hydrolyzed and glucose was detected by addition of 4mL of 0.2% anthrone in H_2_SO_4_ on ice, and subsequent heating at 90°C for 10min. For detection of glucose in conditioned medium, samples were diluted 1/250, 2x the initial volume of 0.2% anthrone in H_2_SO_4_ was added on ice, and samples were subsequently heated at 90°C for 10min. Samples and D-Glucose standards were then transferred to a 96-well plate and the absorbance was measured at 620nm. Glucose concentration in CM was expressed in g/L. Glycogen abundance was inferred from the quantity of glucose extracted from glycogen *per* mg of tissue (μg/mg). Glycogen detection in mouse liver was used as a positive control.

### Interleukin-6 assay

Interleukin-6 (IL6) was measured in CM using a Quantikine mouse IL6 ELISA kit (R&D Systems #M6000B) according to the manufacturer’s instructions. The concentration of IL6 was expressed in pg/mL. Negative values were set to 0.

### Cytokine array

The presence of a range of 40 cytokines and chemokines were investigated in CM using a Proteome Profiler Mouse Cytokine Array kit (R&D Systems #ARY006) according to the manufacturer’s instructions. Membranes were analyzed using Chemidoc Touch and cytokines were quantified on ImageLab software normalized to the technical control.

### Statistical analyses

Experimental data are presented as mean +/- standard error of the mean (SEM). Independent biological replicates are represented in the figures as individual dots, and the total number of replicates (n) *per* group is indicated in the figure legends. Statistical analyses were performed using Prism 11 software (GraphPad). Normality was tested with Shapiro-Wilk’s test and estimated by the visual inspection of Q-Q plots. Statistical tests are indicated in the figure legends. The alpha was set at p≤0.05.

## RESULTS

### Establishment of an ex vivo model to induce muscle contraction by nerve stimulation

To invoke a physiological contraction of the EDL muscle *ex vivo*, starting with acetylcholine (ACh) release at the neuromuscular junction, we sought to induce EDL contraction by direct stimulation of the nerve (Khairullin et al., 2023; Magown et al., 2015). To this end, EDL muscles were dissected with 8-10 mm of the peroneal branch of the sciatic nerve, and mounted in a setup allowing the measurement of contractile force upon electrical stimulation (**Figure S1C-D)**. Given our aim to address interactions between contraction-induced changes in the local muscle environment and cellular responses, we performed subsequent stimulation in the absence of supra-physiological concentrations of oxygen, and in the presence of conventional cell culture media. Muscles were placed at the average *in vivo* length (*i.e.*, 14 mm), which does not differ from the optimal length *ex vivo* (**Figure S2A**). The average optimal stimulation frequency was determined by subjecting the nerve to 600ms pulse trains (10V) at increasing frequencies, demonstrating that stimulation at 25Hz resulted in a fused tetanus (**Figure S2B**), and that maximal force production was reached at 40Hz (**Figure 1A-B**). The average optimal stimulation intensity was determined by subjecting the nerve to 600ms pulse trains (50Hz) at increasing voltages, demonstrating that force production reached a plateau at 5V, and only increased further at 50V (**Figure 1C-D**). Nerve stimulation at supramaximal voltage (10V) resulted in an F_max_ of 25.63 +/- 8.83mN compared to 188.30 +/- 10.81mN when contraction was invoked by field stimulation (50V) (**Figure 1E**). This difference may be partially due to a differential recruitment pattern, as nerve stimulation-induced contraction tended to result in a higher rate of fatigue, and resulted in a greater loss of force at the end of the fatigue protocol than field stimulation-induced contraction (**Figure 1F-H**).

**Figure 1:**
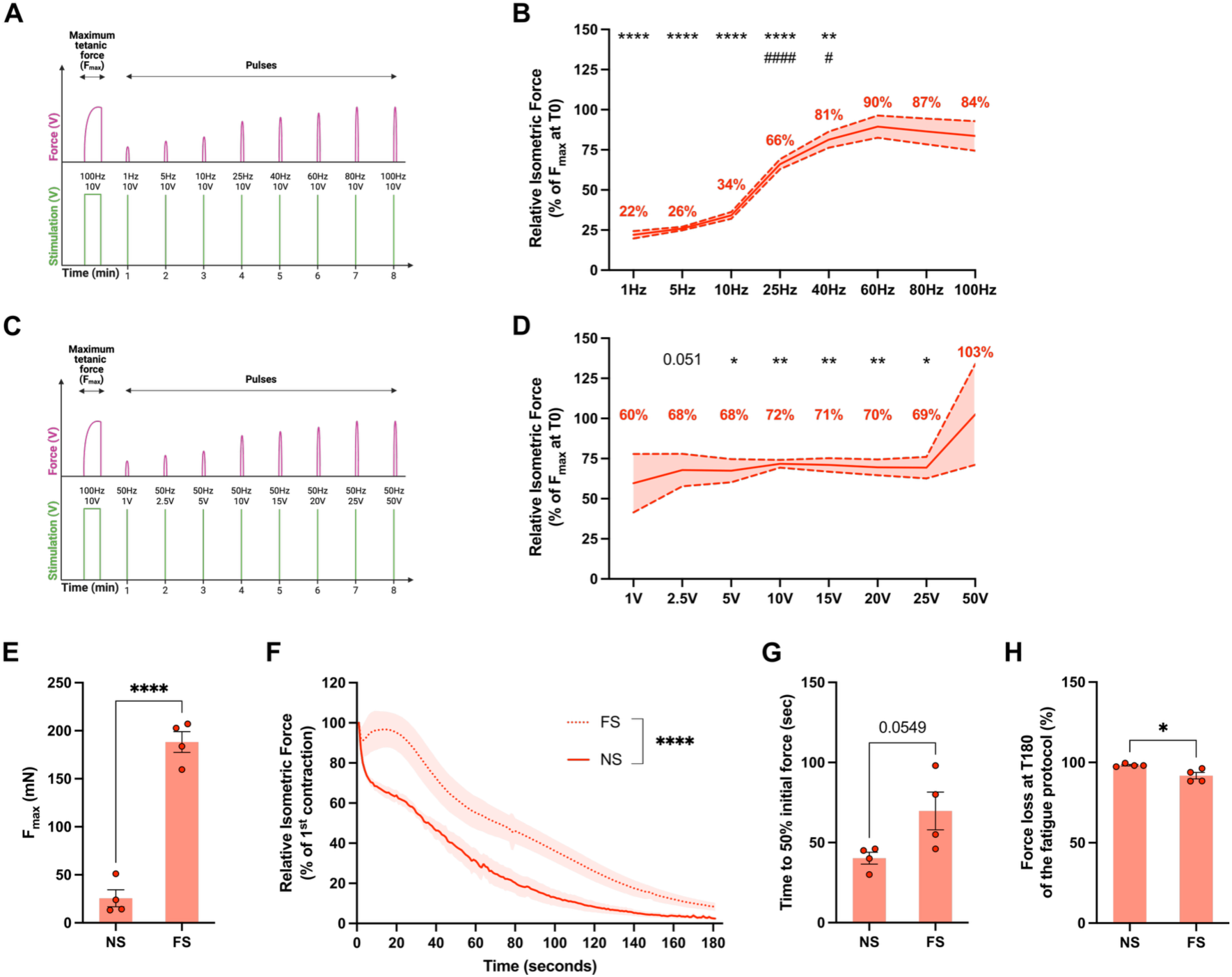
Establishment of an *ex vivo* model to induce muscle contraction by nerve stimulation. **A.** Schematic representation of the force-frequency curve generation. **B.** Muscle force upon increasing stimulation frequency at 10V relative to maximum force (F_max_) at t=0 minutes (min); n=8. **C.** Schematic representation of the force-voltage curve generation. **D.** Muscle force upon increasing stimulation intensity at 50Hz relative to F_max_ at t=0min; n=4. **E.** F_max_ from nerve stimulated (NS) and field stimulated (FS) muscles in mN; n=4. **F.** Muscle force during the fatigue protocol relative to the first contraction at t=0min; n=4. **G.** Time to reach 50% of the initial force in seconds; n=4. **H.** Relative force loss at the end (t=180min) of the fatigue protocol; n=4. Solid lines and bars represent mean +/- SEM. *p<0.05, **p<0.01, ****p<0.0001 compared to t=0min or between indicated bars; ^#^<0.05, ^####^<0.0001 compared to previous timepoint, analyzed using two-way ANOVA with Fisher’s LSD post-hoc test (B, D, F) or unpaired t-test (E, G, H).

### Functional and metabolic response of the EDL muscle to ex vivo contraction by nerve stimulation

Based on the findings above, muscles were placed at the average *in vivo* length, and nerves were stimulated at supramaximal voltage and frequency (10V, 50Hz) during a 1h electrical pulse stimulation (EPS) protocol (**Figure S1E**), which resulted in a 45% decrease of F_max_ at 15min of stimulation, that did not decrease further, yielding a residual force of 43% of F_max_ after 60min of stimulation (**Figure 2A**). Of note, EDL that was not subjected to the EPS protocol demonstrated a progressive increase in F_max_ at 15 and 30 minutes (**Figure S2C**), demonstrating a maintenance of excitation and contractile function over time. Stimulation of contraction *via* the nerve was validated by the removal of the nerve, leading to the absence of force production upon F_max_ measurement (**Figure 2B**). Moreover, curare treatment blocking the nicotinic acetylcholine receptors (AChR) at the neuromuscular junction, resulted in a complete loss of force that was recovered after washout (**Figure 2C**), further confirming stimulation of contraction *via* the nerve.

**Figure 2:**
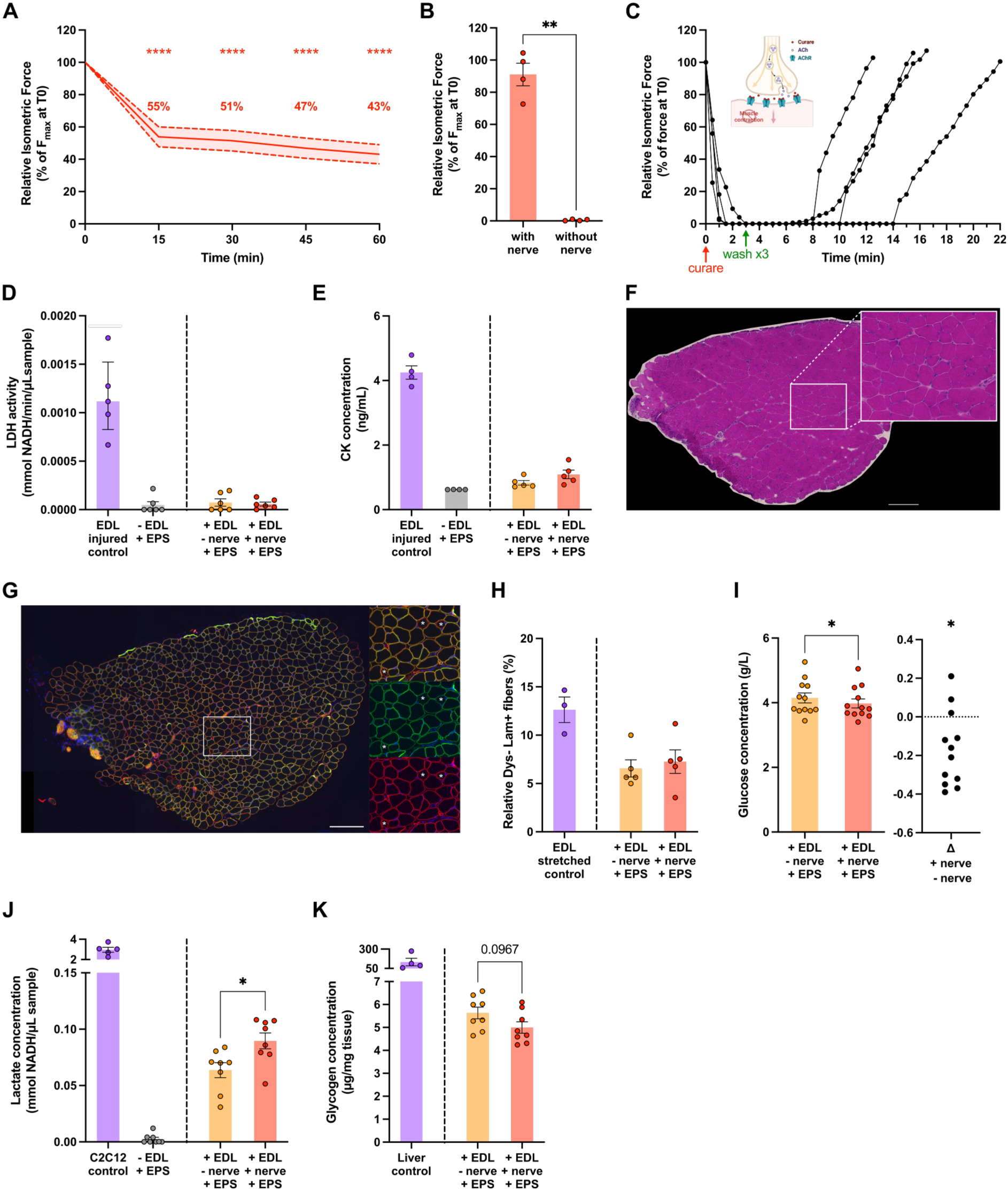
Functional and metabolic response of the *Extensor Digitorum Longus* muscle to *ex vivo* contraction by nerve stimulation. **A.** Muscle maximal force (F_max_) during the electrical pulse stimulation (EPS) protocol relative to F_max_ at t=0 minutes (min); n=11. **B.** Muscle force before and after removal of the nerve relative to F_max_; n=4. **C.** Muscle force during the EPS protocol relative to the first contraction at t=0min; after the first pulse train, muscles were treated with 10µM curare. Three min after treatment, medium was removed and replaced 3 times by fresh medium; n=4. ACh: acetylcholine, AChR: acetylcholine receptor. Lines with dotes represent individual biological replicates. **D.** Lactate dehydrogenase (LDH) activity in conditioned medium (CM) from muscles after EPS protocol; n=6. **E.** Creatine kinase (CK) concentration in CM from muscles after EPS protocol; n=5. **F.** Representative image of H&E staining in EPS-stimulated EDL. **G.** Representative image of dystrophin (Dys, green), laminin (Lam, red), and Hoechst (blue) staining in EPS-stimulated EDL. White asterisks represent Dystrophin negative, Laminin positive (Dys-Lam+) fibers. Scale bars represent 200µm (F,G). **H.** Relative number of Dys-Lam+ fibers compared to the total number of Lam+ fibers, n=3-5. **I.** Glucose concentration in CM from muscles after EPS protocol (left) and differences plot displaying the delta between non-contracted and contracted muscles (right); n=12. **J.** Lactate concentration in CM from muscles after EPS protocol; n=8. **K.** Glycogen concentration in muscles after EPS protocol; n=8. Solid lines and bars represent mean +/- SEM. *p<0.05, **p<0.01, ****p<0.0001 between indicated bars, analyzed using two-way ANOVA with Fisher’s LSD post-hoc test (A), paired t-test (B, I), one-way ANOVA with Fisher’s LSD post-hoc test (H) or unpaired t-test (J-K).

Structural integrity of the EDL after dissection and in response to the EPS protocol was demonstrated by the absence of the cytoplasmic enzymes lactate dehydrogenase (LDH) and creatine kinase (CK) in the conditioned medium (CM) (**Figure 2D-E**). Furthermore, histological analyses demonstrated the absence of sarcoplasmic pallor, immune cell infiltration and centronucleated myofibres (n=3-5) (**Figure 2F**). Moreover, the EPS protocol did not induce any signs of myofiber damage and regeneration (**Figure 2G-H**) (Hildyard et al., 2025).

Repeated muscle contractions are known to increase the metabolic demand of the muscle, that depending on the type and duration of the stimulation results in a depletion of energy stores in the muscle, and the release of metabolic (by)products (Domin et al., 2021; Egan & Zierath, 2013; Hargreaves, 2020). In line, the EPS protocol resulted in a mild but significant decrease in the glucose concentration in the CM (**Figure 2I**), suggesting that EPS stimulates skeletal muscle glucose uptake. Furthermore, lactate was detected in CM from muscle, and its concentration was increased by 41% in CM from EPS-stimulated muscle (**Figure 2J**), suggesting that EPS increased lactate release into the CM. In line, EDL glycogen content tended to decrease by 11% in EPS-stimulated muscle (**Figure 2K**). Altogether, this suggests a metabolic shift towards glycolysis upon the EPS protocol, which evokes physiologically relevant metabolic alterations to the local skeletal muscle environment.

### Functional and metabolic recovery of the EDL muscle after cessation of ex vivo contraction by nerve stimulation

Skeletal muscle metabolism and function is not just altered acutely during muscle contractions, but is also modulated during the hours following cessation of contraction (Ato et al., 2017; Baker et al., 1993; Louis et al., 2007; Raastad & Hallén, 2000; Sahlin & Ren, 1989; Sayers & Clarkson, 2001). To assess if *ex vivo* EDL can functionally recover after the EPS protocol, F_max_ was measured at 1h or 3h after cessation of the EPS protocol. At 1h post-EPS, muscle force tended to be increased as compared to directly after the EPS protocol, returning to 91% of initial F_max_ (**Figure 3A**), whereas no force recovery was detected at 3h post-EPS (**Figure 3B**).

**Figure 3:**
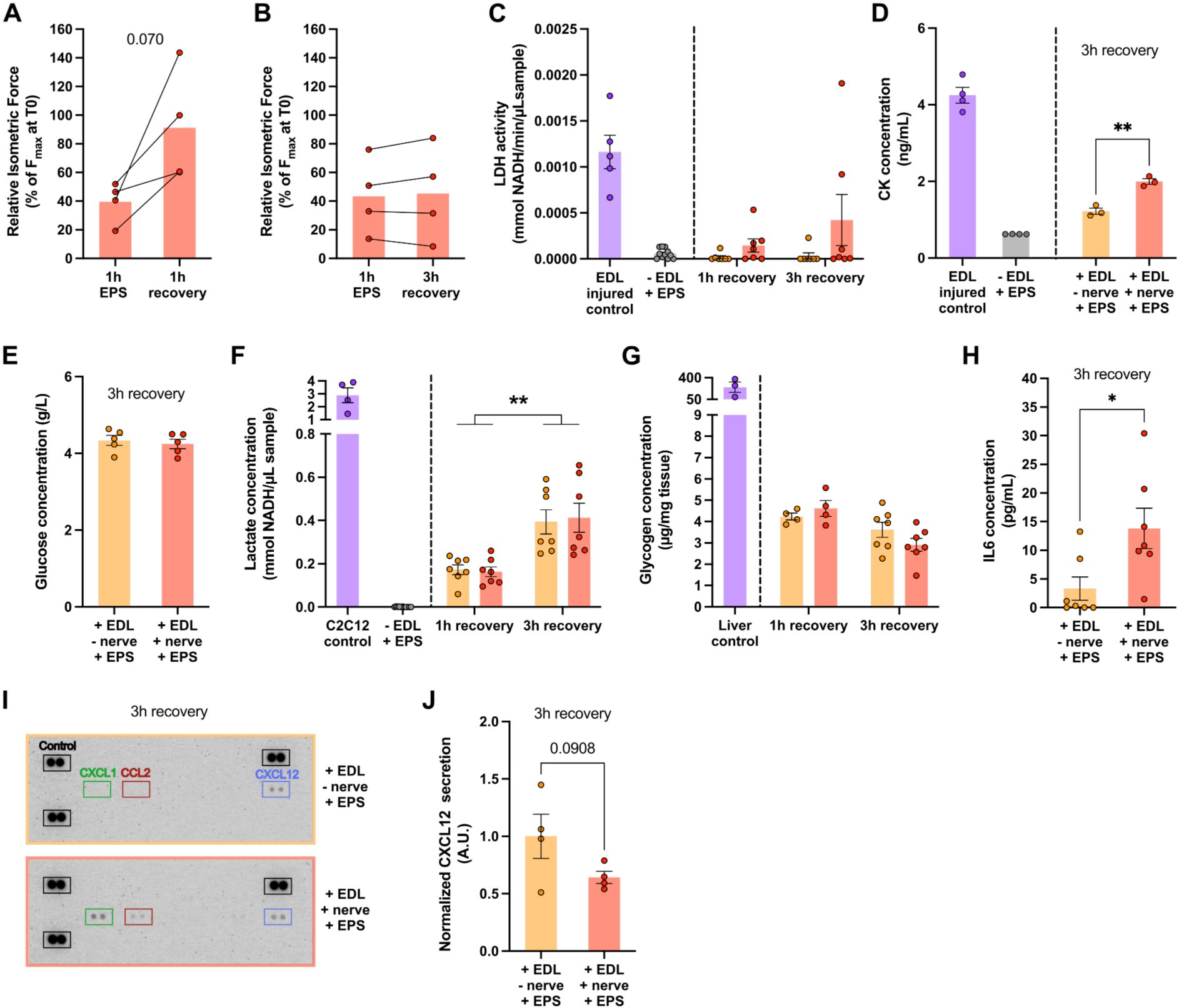
Functional and metabolic recovery of the *Extensor Digitorum Longus* muscle after cessation of *ex vivo* contraction by nerve stimulation. **A-B.** Muscle maximal force (F_max_) relative to F_max_ at t=0 minutes (min), immediately after electrical pulse stimulation (EPS) protocol and at 1 hour (h) (A) or 3h (B) recovery; n=4. **C.** Lactate dehydrogenase (LDH) activity in conditioned medium (CM) from muscles at 1h and 3h recovery; n=7. **D.** Creatine kinase (CK) concentration in CM from muscles at 1h and 3h recovery; n=3. **E.** Glucose concentration in CM from muscles at 3h recovery; n=5. **F.** Lactate concentration in CM from muscles at 1h and 3h recovery; n=7. **G.** Glycogen concentration in muscles at 1h and 3h recovery; n=4-7. **H.** Interleukin-6 (IL6) concentration in CM from muscles at 3h recovery; n=7. **I.** Representative image of cytokine array of CM from muscles at 3h recovery. CXCL: C-X-C motif chemokine ligand. CCL: C-C motif chemokine ligand. **J.** Quantification of CXCL12 from cytokine array of CM from muscles at 3h recovery; n=4. Bars represent mean +/- SEM. *p<0.05, **p<0.01 between indicated bars, analyzed using test paired t-test (A, J), unpaired t-test (D, H) or two-way ANOVA with Fisher’s LSD post-hoc test (F-G).

Structural integrity of the EDL during the 1h post-EPS was demonstrated by the absence of LDH in the CM (**Figure 3C**). At 3h post-EPS LDH was detected in 2/7 muscles, and CK was increased in CM from EDL that were prior subjected to the EPS protocol (**Figure 3C-D**), suggesting a late-onset cytoplasmic release after contraction.

At 3h post-EPS, no difference in glucose concentration was observed, indicating that glucose may no longer be the primary energy substrate utilized by the muscles at this time point (**Figure 3E**). Lactate was detected in CM from muscle at 1h and 3h post-EPS, but its concentration was not altered upon prior EPS (**Figure 3F**), suggesting an increase in lactate re-uptake in EPS-stimulated muscle during the first hour after cessation of EPS (**Figures 2J & 3F**). In line, at 1h and 3h post-EPS, EDL glycogen content did not differ between stimulated and non-stimulated muscles (**Figure 3G**), which suggests a recovery of glycogen stores.

Skeletal muscle contraction is not only associated with altered substrate use and metabolism, but also results in the release of myokines (a.k.a. exerkines) (Hoffmann & Weigert, 2017; Severinsen & Pedersen, 2020). The myokine IL6 is known to be released after acute exercise (Petersen & Pedersen, 2005), and while non-detectable at 1h post-EPS, IL6 was detected in CM at 3h post-EPS and was strongly increased in CM from stimulated compared to non-stimulated muscle (*i.e.* 4.2-fold; **Figure 3H**). Furthermore, a cytokine array on CM at the 3h post-EPS timepoint demonstrated the detection of CXCL1, CCL2/MCP-1, and CXCL12 (**Figure 3I**). Interestingly, CXCL12 abundance at 3h post-EPS was lower in CM from contracted than non-contracted muscle (**Figure 3J**).

### Functional and metabolic response of the EDL muscle to ex vivo contraction induced by field stimulation

While the physiological relevance of muscle contraction induced *via* nerve stimulation is evident, the relatively low force production upon F_max_ induction may limit EPS-induced metabolic alterations. To address this, we subjected EDL muscles to *ex vivo* contraction induced by field stimulation (FS) (**Figure S3A-C**). The EPS protocol by FS resulted in a loss of 70% of F_max_ at 15min, and a progressive loss of force for the rest of the protocol, resulting in 3% of F_max_ at 60 min (**Figure S3D**), which is a similar if not milder force loss compared to nerve stimulation (NS). Structural integrity of the EDL in response to the EPS protocol induced by FS was demonstrated by the absence of lactate dehydrogenase (LDH) in the CM (**Figure S3E**). While the higher absolute force produced upon FS than upon NS may imply a higher metabolic demand (**Figure 1E**), the EPS protocol by FS induced lactate release in the CM by 33%, which is similar to the induction by NS (**Figure S3F**). Yet, the depletion of glycogen stores by 58% in stimulated *versus* non-stimulated muscle was more robust than upon the induction by NS (**Figure S3G**). Functional recovery was not observed at either 1h or 3h post-EPS (**Figure S3H-I**), which demonstrates the supraphysiological effects of the EPS protocol when induced by field stimulation.

## DISCUSSION

Skeletal muscle contractile and metabolic functions are coordinated through muscle-derived endocrine, paracrine and autocrine signaling. To address such signals in a physiologically relevant manner, we developed a setup of nerve-induced *ex vivo* isometric contraction of whole muscle. This model initiates muscle contraction starting with the release of ACh at the neuromuscular junction, and allows the collection of released and depleted factors at multiple timepoints for identification and quantification.

Myofiber-derived paracrine signaling has previously been explored using conditioned medium (CM) from *in vitro* differentiating myoblasts (Henningsen et al., 2010; Le Bihan et al., 2012), or from *in vitro* differentiated myoblasts after contraction induced by electrical pulse stimulation (EPS) (Bydak et al., 2022; Raschke et al., 2013; Scheler et al., 2015). However, these *in vitro* models suffer from an immature contractile and metabolic system, and completely exclude the effect of the muscle architecture and other cell types. The complex interplay between cell-types and structural components through paracrine signaling has been explored by unbiased approaches. McKellar *et al*. performed computational ligand-receptor analysis of integrated skeletal muscle single-cell and single-nucleus transcriptomics data to predict cellular interactions through ‘secreted signaling’ (McKellar et al., 2021). Furthermore, a recent study profiled the skeletal muscle secretome in extracellular fluid collected by low-speed centrifugation (Mittenbühler et al., 2023). While valuable, these approaches do not directly address the effect of increased muscle contractility, nor the release and extraction of local factors such as myokines and metabolites. In addition, extracellular fluids are collected for endpoint measurement(s), and therefore do not allow assessment of the kinetics of paracrine signals.

We demonstrated that nerve-induced *ex vivo* skeletal muscle contraction results in a mild but consistent force production. Maximal force was substantially higher upon field stimulation than upon nerve stimulation, whose absolute force values were in line with previous literature (*e.g.*, Mader et al., 2022). This may reflect a supraphysiological recruitment of myofibers upon field stimulation-induced contraction. Indeed, previous studies suggest that the parameters of stimulation impact motor unit recruitment, and thereby both the number and type of contracted myofibers (Bickel et al., 2011; Gorman & Mortimer, 1983; Gregory & Bickel, 2005; Gundersen, 1998). Moreover, in our setup, the muscle is not perfused and relies on passive diffusion of oxygen and nutrients. *Ex vivo* muscle contraction is often executed in a hyperoxic environment (Balasubramanian et al., 2022; Franco et al., 2014; Katz, 2022; Liu et al., 2020; Mendias et al., 2006; Merry et al., 2010; Plant et al., 2001) and the atmospheric conditions used here may lead to a weaker contraction. As opposed to these previous studies characterizing maximal muscle force, here we aimed to study the metabolic response to contraction in a physiologically relevant environment. Indeed, under physiological conditions, *in vivo* skeletal muscle is exposed to an even lower oxygen tension, ranging from 7.5 to 31 mmHg depending on the muscle type and the rate of contraction (Ortiz-Prado et al., 2019; Richardson et al., 2006). Thus, our use of atmospheric conditions better resembles the physiological muscle environment than conventional hyperoxic conditions, and could be even better represented by placing the setup in a hypoxia incubator. In line with a rapid drop of force during the fatigue protocol, the main loss of force during the 1-hour EPS protocol occurred in the first 15 minutes. Interestingly, muscles only subjected to F_max_ demonstrated an increase in force over the first 30 minutes, which may suggest an underestimation of EPS-induced relative force loss. Previous studies also describe such ‘potentiation’ (Łochyński et al., 2021; Zhi et al., 2005), which may be due to a calcium, potassium and nutrient rich environment *ex vivo* (Angelidis et al., 2024; Łochyński et al., 2021; Zhi et al., 2005). Physiological relevant nerve-induced stimulation of the EDL was supported by the full recovery of force production within 1 hour after the EPS protocol, which is in contrast to the lack of force recovery after field stimulation-induced contraction. Nerve-induced contraction resulted in a lower glucose and higher lactate abundance in CM, suggesting increased glucose uptake and lactate release, along with a tendency towards a depletion of muscle glycogen content. These metabolic changes resemble the response of skeletal muscle to repeated contractions *in vivo* (Hearris et al., 2018; van Hall, 2010). The glucose and lactate levels in CM were normalized, and glycogen content returned to similar levels in contracted muscle compared to non-contracted muscle after 1 and 3 hours of recovery, demonstrating an active recovery of the intracellular metabolic status. Interestingly, CK concentration was increased in CM from contracted *versus* non-contracted muscle at 3 hours of recovery. While CK is classically used as a marker of myofiber damage, CK release may also be part of the normal physiological response to muscle contraction (Baird et al., 2012; Baltusnikas et al., 2015; Brancaccio et al., 2007). Altogether, this demonstrates the physiological relevance of our nerve-induced *ex vivo* skeletal muscle contraction method from a contractile and metabolic perspective.

In line with the known *in vivo* physiological response to exercise, IL6 release was induced 3 hours after cessation of the EPS protocol (Chowdhury et al., 2020; Pedersen et al., 2003; Steensberg et al., 2000). Conversely, CXCL12 levels in CM at 3h recovery tended to be lower in contracted compared to non-contracted conditions, suggesting either a decrease of CXCL12 release or an increase in CXCL12 scavenging (Hunger et al., 2012). This seems in contrast to previous studies, which demonstrated increased mRNA and protein expression of CXCL12 and its receptors upon training, as well as exercise-mimicking physiological effects of CXCL12 signaling *in vitro* in C2C12 cultures (Puchert et al., 2016; Yamada et al., 2019). Yet, to our knowledge, no previous studies profiled the full kinetic of CXCL12 release during and after (non-damaging) contraction *in vivo* or *ex vivo*. These contraction-induced alterations in IL6 and CXCL12 in CM provide a proof-of-concept for the use of our model to address contraction-induced muscle-specific paracrine and endocrine factors.

### Limitations of the model

Our model overcomes several limitations associated with previously used methods to study myofiber-derived paracrine signaling, particularly by including the use whole skeletal muscle *ex vivo* to consider the cellular complexity in its response to contraction. While we demonstrated that nerve stimulation leads to a physiologically relevant contraction, one may opt to use field stimulation-induced contraction *e.g.*, to assess acute effects given its high force production. However, to address paracrine signaling, nerve-stimulation benefits from the exclusion of potential effects of the electrical current on the medium or ‘co-cultured’ cells. Indeed, field stimulation could result in electrolysis of the medium, and has been shown to induce ROS generation (Derave et al., 2006). Moreover, electrical currents have been shown to directly affect survival, proliferation and migration of different cell types (Meng et al., 2022; Robinson, 1985; Taghian et al., 2015).

Due to the reliance on passive diffusion, the availability of nutrients in our setup may be suboptimal. *Ex vivo* contraction is classically performed in poor media (*i.e.* Ringer solution (Merry et al., 2010), Tyrode solution (Balasubramanian et al., 2022; Katz, 2022) or Krebs-Ringer-Henseleit buffer (Liu et al., 2016; Mendias et al., 2006; Plant et al., 2001)). To at least partially compensate for decreased nutrient delivery, we use a nutrient-rich medium. However, we still observed an increase in lactate in the CM independent of contraction, which may suggest that the demand for ATP and/or oxygen exceeds supply. Although suboptimal nutrient availability may be improved by increasing the medium volume, here we purposefully kept the medium volume low (*i.e.*, 6mL) to support better detection of paracrine and endocrine signals in CM.

Here, we developed an EPS protocol, that leads to substantial contractile, metabolic and paracrine alterations in EDL. Nevertheless, it should be taken into consideration that the duration of pulse trains, the delay between pulse trains, the total duration of the contraction protocol, and the contraction type (isometric/concentric) all influence the contractile, metabolic and paracrine responses. Moreover, these responses may differ depending on the contractile and metabolic subtype of the skeletal muscle, *e.g.*, when using a more fatigue resistant muscle such as *Soleus* (Schiaffino & Reggiani, 2011).

Taken together, we have developed a physiologically relevant model of nerve-induced *ex vivo* whole muscle contraction, which will help unravel how skeletal muscle mass and function are maintained *via* the complex interplay between cell-types and structural components through paracrine and endocrine signaling upon muscle contraction.

## Supporting information

Video S1. Maximal ex vivo contraction

## AUTHOR CONTRIBUTIONS

RM designed the study, CD, EB, AK and RM conceived experiments, CD, IB and AM performed and analyzed experiments, LL provided technical support, BA provided conceptual input, CD prepared the figures, CD and AK wrote and revised the manuscript, and EB and RM revised the manuscript. All authors read and approved the manuscript.

## SUPPLEMENTAL MATERIALS AND METHODS

### Ex vivo field stimulation-induced muscle contraction setup

The field stimulation (FS) setup is composed of 2 solid carbon electrodes and two surgical steel pillars embedded into resin (Soloplast Epoxy Resin) serving as a customized lid fitting a standard 6-well culture plate (**Figure S3A-B**). This lid holds and stimulates up to six muscles at the same time. *Extensor Digitorum Longus* (EDL) muscles were mounted at 14mm (*i.e. in vivo* and optimal length) submerged in pre-warmed (37°C) DMEM high glucose GlutaMAX medium (Gibco #31966021) supplemented with 1% penicillin-streptomycin (Sigma #P4333). Mounted EDL muscles were then acclimatized in a humidified incubator at 37°C and 5% CO_2_ for a 15-minute (min) period before subsequent force measurements or the electrical pulse stimulation (EPS) protocol.

To assess the preservation of force in unstimulated muscles, the non-contracting controls were subjected to F_max_ measurements at the start, and after each 15min interval. For measurements of metabolic responses in muscles and conditioned media (CM) upon nerve-stimulation, non-contracting controls received no electrical stimulation, resulting in no contraction. Moreover, an EPS control CM was generated by subjecting the EPS protocol stimulation to medium in the absence of an EDL (**Figure S3C**).

## SUPPLEMENTAL FIGURES

**Supplemental figure 1:**
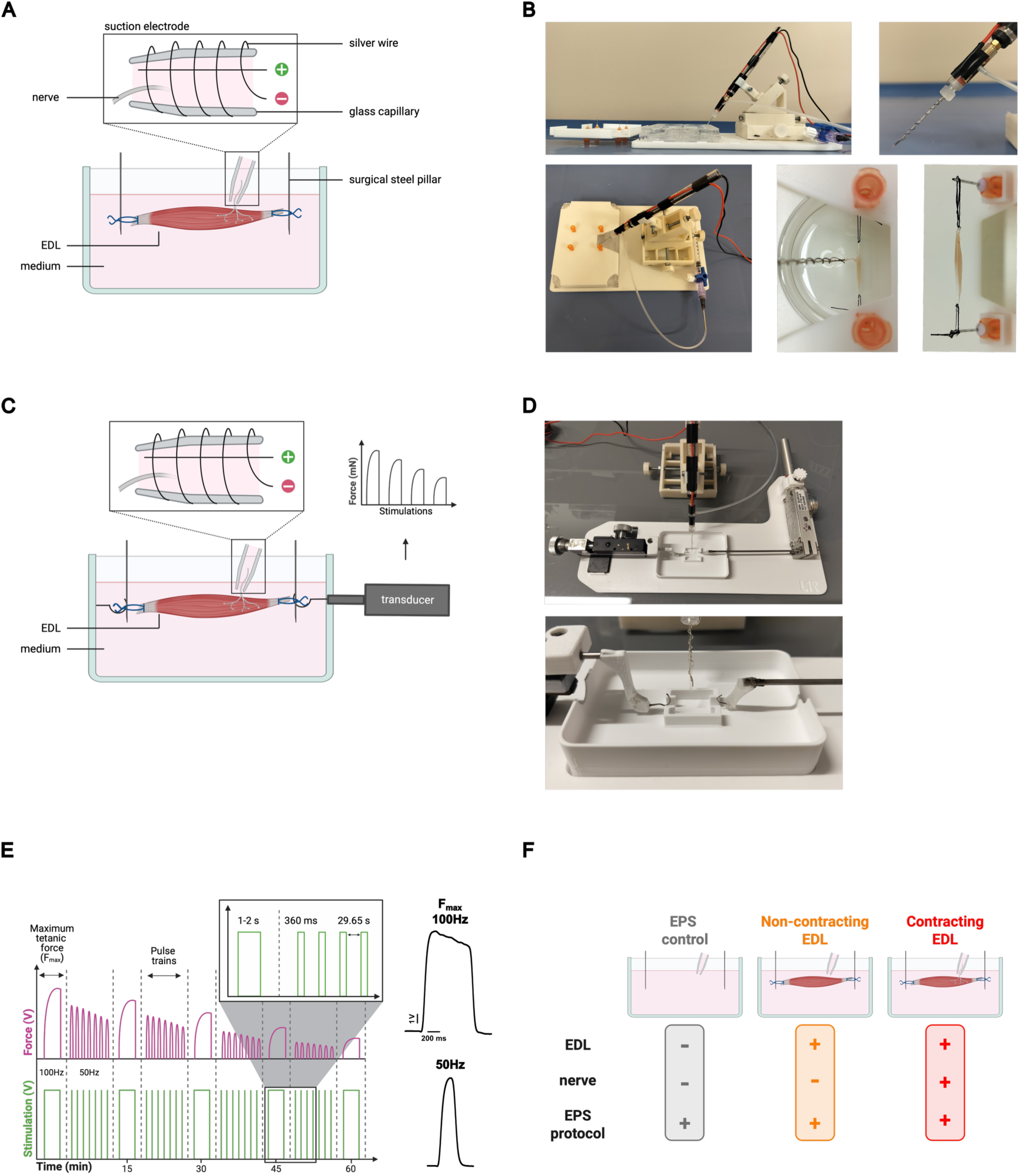
Establishment of an *ex vivo* model to induce *Extensor Digitorum Longus* muscle contraction by nerve stimulation. **A.** Schematic representation of nerve stimulation (NS) setup. **B.** Images of NS setup; side view (top left); zoom of suction electrode (top right); mounted *Extensor Digitorum Longus* (EDL) in setup top view (bottom left); EDL muscle with nerve aspirated into the suction electrode (middle); EDL mounted at *in situ* length (bottom right). **C.** Schematic representation of the force measurement setup with force-displacement transducer. **D.** Images of force measurement setup. **E.** Schematic representation of the electrical pulse stimulation (EPS) protocol, and representative traces of maximum force (F_max_,1-2sec, 10V, 100Hz) and pulse train (360 milliseconds (ms), 10V, 50Hz). **F.** Schematic representation of the experimental conditions in the NS setup.

**Supplemental figure 2:**
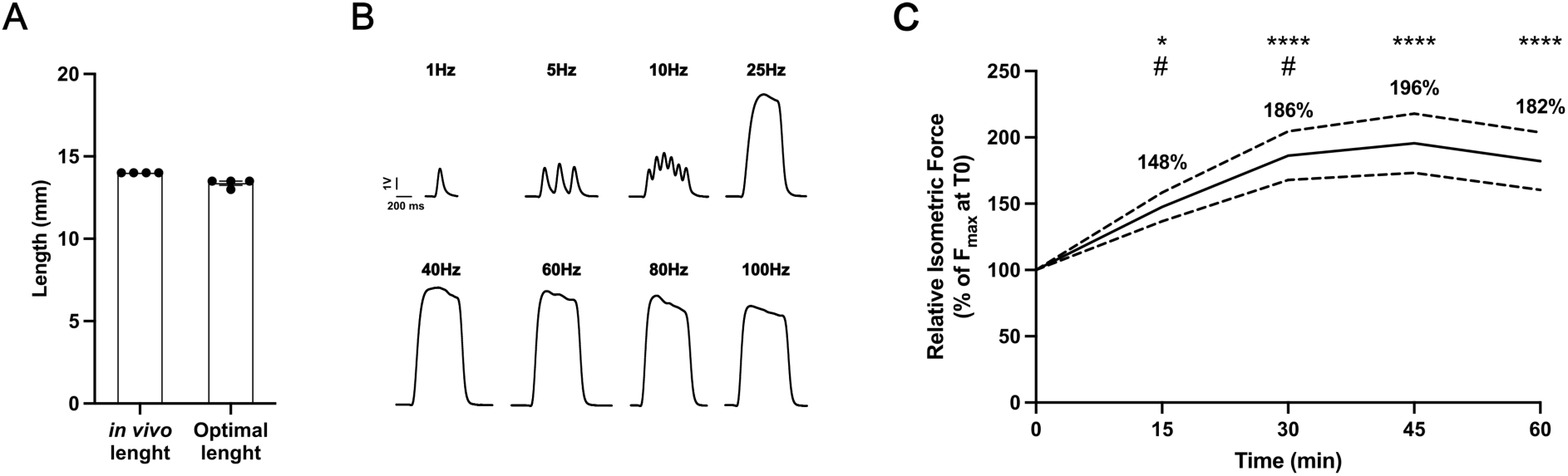
Functional response of the *Extensor Digitorum Longus* muscle to *ex vivo* contraction by nerve stimulation. **A.** In vivo and optimal muscle length of *Extensor Digitorum Longus* (EDL). **B.** Representative traces of pulse trains at indicated frequencies (600 milliseconds (ms), 10V), corresponding. **C.** Relative isometric force relative to muscle maximal force (F_max_) at t=0 minutes (min) in muscles not subjected to electrical pulse stimulation (EPS); n=8. Solid lines and bars represent mean +/- SEM. *p<0.01, ****p<0.0001 compared to t=0min, ^#^p<0.05 compared to previous timepoint, analyzed by two-way ANOVA with Fisher’s LSD post-hoc test.

**Supplemental figure 3:**
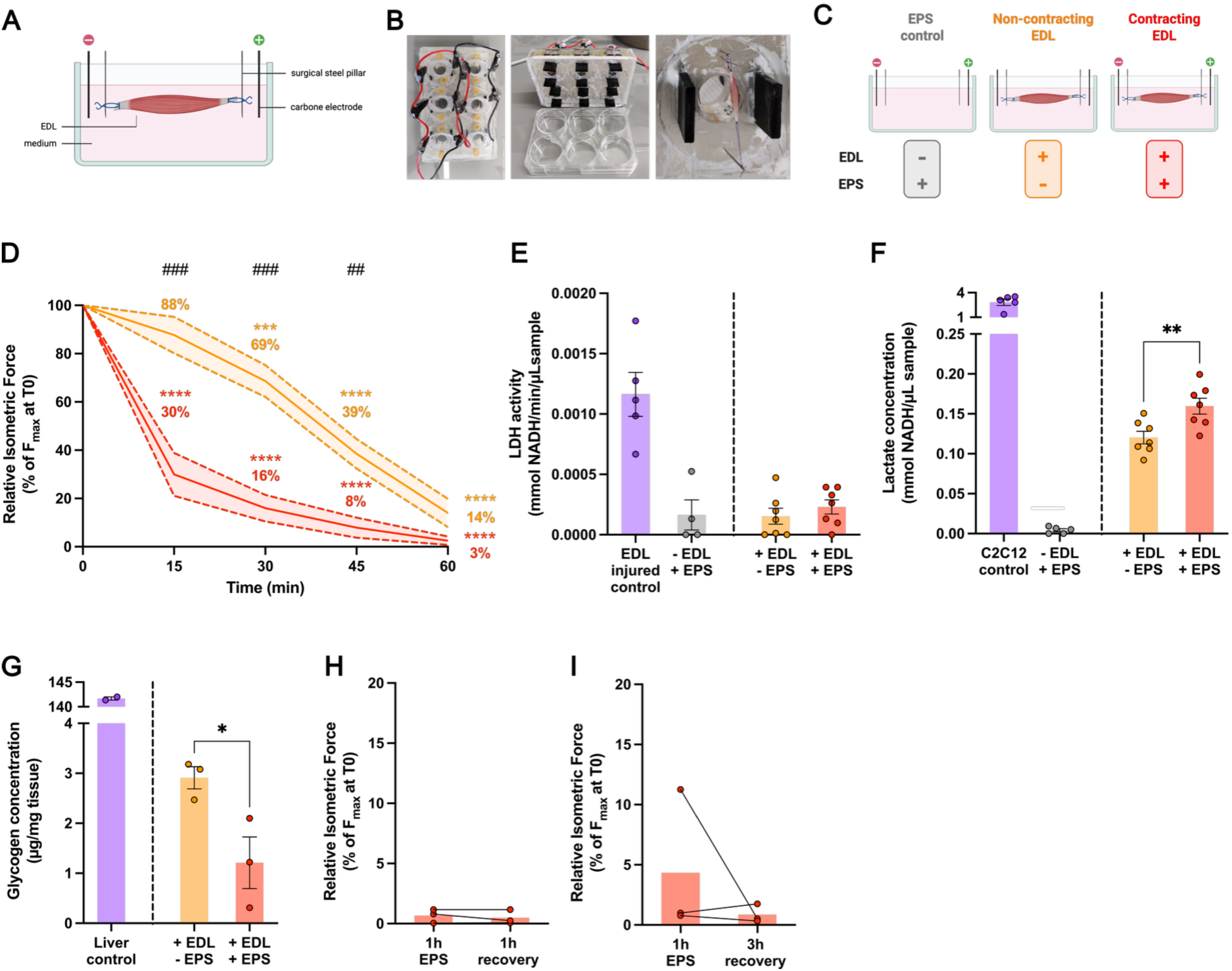
Functional and metabolic response of the *Extensor Digitorum Longus* muscle to *ex vivo* contraction by field stimulation. **A.** Schematic representation of field stimulation (FS) setup. **B.** Images of FS setup; top view (left); inside view with lid up (middle); isolated *Extensor Digitorum Longus* (EDL) muscle (right). **C.** Schematic representation of the experimental conditions in FS. **D.** Muscle maximal force (F_max_) during the electrical pulse stimulation (EPS) protocol relative to F_max_ at t=0 minutes (min); n=6; * compared to t=0min and ^#^ compared to non-stimulated control. **E.** Lactate dehydrogenase (LDH) activity in conditioned medium (CM) from muscles after EPS protocol; n=7. **F.** Lactate concentration in CM from muscles in CM from muscles after EPS protocol; n=7. **G.** Glycogen concentration in muscles after EPS protocol; n=3. **H-I.** F_max_ relative to F_max_ at t=0min, immediately after EPS protocol, and at 1 hour (h) (H) or 3h (I) recovery. Solid lines and bars represent mean +/- SEM. *p<0.05, **/^##^p<0.01, ***/^###^p<0.001, ****p<0.0001 between indicated bars, analyzed using two-way ANOVA with Fisher’s LSD post-hoc test (D) or unpaired t-test (F, G).

## SUPPLEMENTAL VIDEO

**Video S1: Maximal *ex vivo* contraction of *Extensor Digitorum Longus* muscle by nerve stimulation**

